# Comparative assessment of morphology, anatomy and phylogeny of two exceptionally rare narrow endemic *Tephrosia spp*. (Leguminosae) from the Indian subcontinent

**DOI:** 10.1101/2022.08.14.503883

**Authors:** Akash Vanzara, Gagandeep Kaur Bhambra, Padamnabhi S. Nagar

## Abstract

Leguminosae is the third most abundant family of angiosperm in the world, and have a distinct difference that serves to identify and differentiate closely related species. Among 650 genera and 18,000 species of legumes in India, *Tephrosia* is the most emerged genus, of which two endangered and narrow endemic species *Tephrosia jamnagarensis* Sant and *T. colina* var. *lanuginocarpa* Sharma, possess a greater challenges for biological, ecological, and competitive improvement than other plants. Both the species are very unique and only found in a few isolated pockets of Gujarat around the globe. Therefore, in light of this, the present research aims to understand how morphological, anatomical, and genetically both species vary based on their variability nature are contextualized in an Indian context. Studies carried out on Morpho-anatomical observations and classical taxonomic approaches, such as the axillary cyme vs terminal raceme in *T. jamnagarensis* vs *T. colina* var. *lanuginocarpa* Sharma, the corolla being 4.5 cm long vs 1.5 cm long, the wings being 5.5 cm long vs 1.40 cm long, the keel being 3.4 cm long vs 1.5 cm long. First, ever genetic identity of the plants was assessed by using *matK* gene of the *rbcl* chloroplast region, and phylogenetic trees were constructed using sister clade separation. Sequences of both the plant species and others species that are found in India were submitted to NCBI and the embryonic stage of seeds were then used to examine seed morphology and propagation.

## Introduction

As one of the major groups of Angiosperms (flowering plants), *Tephrosia* encompasses more than 350 species of woody annual and perennial herbs, which are widely distributed throughout tropical and subtropical regions of the world (Samuel, Mahesh, and Murugan 2019). The name of *Tephrosia* was accepted at the International Botanical Congress in 1905, and nomenclature was explained (Wood 1949), but it was previously merged with *Needhamia* Scop. (1777) and *Reineria Moench* (1802) as *nomina rejecienda. Cracca* L. (1753) and continue recognized as *Cracca* L. by adherents of American code during 1930s. In Flora Zeylandica (1747), Linnaeus referred to the plant as *Cracca*, but Candolle in (1826) endorsed its maintenance of name. (Phillip, 1986) asserted French word means “staying green.” Whereas (Quattrocchi, U. 2017) the Greek word (“Tephros”) means “ash-colored”. Currently, there are thirty species and one variety present in South Asia (Kumar P.V, 2003), of which twenty seven species and one variety have been documented for India (Sanjappa 1992), while 10 species and one subspecies have been discovered in Gujarat (Shah 1978). The type locality for *Tephrosia jamnagarensis* is jamnagar, the state of its discovery (Shah 1978),(Thaker 1910). Species of *Jamnagarensis* have a specific epithet describing their type location. A wide variety of medicinal uses of *Tephrosia* genus have been found, including treatments for diabetes, ulcers, diarrhea, wound healing, anti-inflammatory, insecticidal, antiviral, anti-protozoal, anti-fungal, anti-plasmodial, and many others (Chen et al. 2014). There are five main classes of compounds found in Tephrosia, including flavonoids, rotenoids, terpenoids, sterols, essential oils, and fixed oils. The most common compound isolated and identified was flavonoids. Stereochemistry has been studied for some compounds, for example, Praecansone from Tephrosia pumila exists as two isomers (Dagne, Yenesew, and Waterman 1989).

Based on several studies conducted by the taxonomists, *Tephrosia* was divided into four sections *viz*. *Mundulea, Brissonia, Craccoides, and Reineria*. Of which *Mudulea* and *Reineria* were predominant in India. As the genus evolved, it was further classified into three subgenera including Marconyx (which includes *T. tenuis*), Brissonia (which includes *T. candida*), and Reineria (which includes the remaining *Tephrosia* species) (Lakshmi et al. 2008). A significant feature of this genus is its taxonomic complexity (G. P. Lewis et al. 2005). In light of all; identification of these species is a vital and challenging task. The anatomical information provided by plants also proves to be useful for taxonomic identification at all levels of the taxonomic hierarchy, as well as for assessing taxonomic relationships among the other taxa (Stuessy, Crawford, and Marticorena 1990).

For systematic purposes, plant morphology largely serves as a tool for carving diversity into its systematic components based on morphological characteristics (Kaplan 2001). Although a comprehensive phylogeny of the genus has yet to be determined. Therefore, present investigation attempts to provide a consistent and reliable diagnosis for the two narrow endemic species *Tephrosia jamnagarensis* Sant and *T. Colina* var *Lanuginocarpa* Sharma, Thus method can be extremely useful for delimitation of the species even in fragmentary circumstances based on taxonomy, anatomy, and phylogeny.

## Materials and Methodology

### Plant collection and Identification

The investigated species of *Tephrosia* were cultivated in the arboretum of the M.S. University in Vadodara, Gujarat. All fully grown plant part (root, stem, leaves and seeds) were collected. The morphological characteristics of the species were examined under the Leica Microsystems S9E Stereo microscope and key characters of genus and species were identified with the help of relevant literatures, including state as well as national floras (Cook n.d.), (Bole J.M. 1988), (Shah 1978), (Karthikeyan Sanjappa, M. & Moorthy, S. 2009).

### Extraction of DNA and PCR

Fresh leaf samples weighing 50 mg were crushed in liquid nitrogen and fine powder was obtained by crushing them in a sterile mortar and pestle. The DNA extraction was carried out using Plant/Fungi DNA isolation kit (Sigma Aldrich Cat# E5038). The PCR reaction was conducted in a volume of 20μl containing 1X DreamTaq Green PCR Master Mix (Cat# K1081), 50 ng of genomic DNA, and 10 pmol of primers. The following conditions were used for PCR reactions: 94°C for 4 min, followed by 35 cycles of 94°C for 30 sec, annealing at 56°C for 30 sec, and extension at 72°C for 1 minute. matK & rbcL primer (Yu, XUE, and ZHOU 2011), was used for amplification of chloroplast DNA sequences at 72°C for 10 minutes and stored at 4°C after the final extension. The purification and sequencing of the amplified PCR product was performed at Gujarat State Biotechnology Mission, using Purelink TM Quick PCR Purification Kit (Cat#K310001).

### Analysis of sequence data based on phylogenetic relationships

In order to conduct phylogenetic analysis, we downloaded 23 sequences of *Tephrosia* of which 12 sequences were of rbcL gene and 11 sequences of matK gene from GenBank in FASTA format (Table.5). In Bioedit 7.2.5 (Hall 1999), ambiguities were resolved by aligning forward and reverse sequences. A nucleotide sequence alignment was performed with ClustalW (Thompson, Gibson, and Higgins 2003). Embedded into MEGA 6.0 (Tamura et al. 2013). A phylogenetic analysis was conducted on the concatenated dataset using Partition Finder (Lanfear et al. 2012) to choose the best partitioning scheme. A partition scheme was chosen for both ML analyses and phylogenetic inferences using RaxML (Silvestro and Michalak 2012). Based on 1000 bootstrap replicates with a GTR + I model, clade support was assessed

### Anatomical preparations

Collected samples were fixated with FAA fixation system (Berlyn 1976). After following fixation protocol, samples were washed with water and further processed for Tertiary Butyl Alcohol (TBA) series before being permeated within paraffin wax. The sections were cut in the plains at 15-20 μm thicknesses and stained with Safranin (0.5% with water) and mounted on 50% glycerin. The obtained sections were examined under Lecia DM 2000 microscope and photographs were captured from digital canon camera connected to the microscope.

### Seed germination and phenology

The morphology and variability in mature fruit and seed was examined under the microscope. The length and diameter was measured with the help of electronic scale in (Software name). In order to study the germination of seeds 36N H_2_SO_4_ was applied for two minutes. Thereafter, seeds were washed thoroughly to remove acid residual traces under tap water. Similar experiments were carried out with hot water, cold water and with various concentrations of GA_3_ (Gibberellic acid) (250 ppm, 500 ppm)

## Result

A description of the observed morphological key characteristics of *T. jamnagarensis* and *T. collina* is given in Tables 1 and illustrated in Figures 1.

**Table.1:** Comparative morphology of *T. jamnagarensis* and its allied species, *T. collina var lanuginocarpa*

|  | Part of the plant | Name of the plant |  |
| --- | --- | --- | --- |
|  |  | *T. jamnagarensis* | *T. collina var lanuginocarpa* |
| Morphology of the Plant | Habitat | Annual herb with 60-82 cm long | Annual herb with 60-100 cm long |
|  | Stem | Simple, densely appressed silky hairs erect to sub erect cylindrical | Simple thinly pubescent towards base angled |
|  | Leaves | Simple, stipules subulate, up to 3mm long, uninerved, pubescent on margin<br>lamina subacute-obtuse at apex, rarely mucronate, adaxially sub glabrous (Fig ) and silky villous abaxially (Fig ), | Compound imparipinnate stipules up to 8mm long, subulate, 3 nerved persistent, lamina obtuse or somewhat truncately emarginate at apex, rarely mucronate, sericeous abaxially and adaxially glabrous |
|  | Inflorescence | Axillary cyme | Terminal raceme |
|  | Flower | Flower bracteates, ebracteolate | Bract filiform, bracteolate |
|  | Gynoceium | 4 mm subsessile | 1.3 mm sessile |

**Fig.1.**
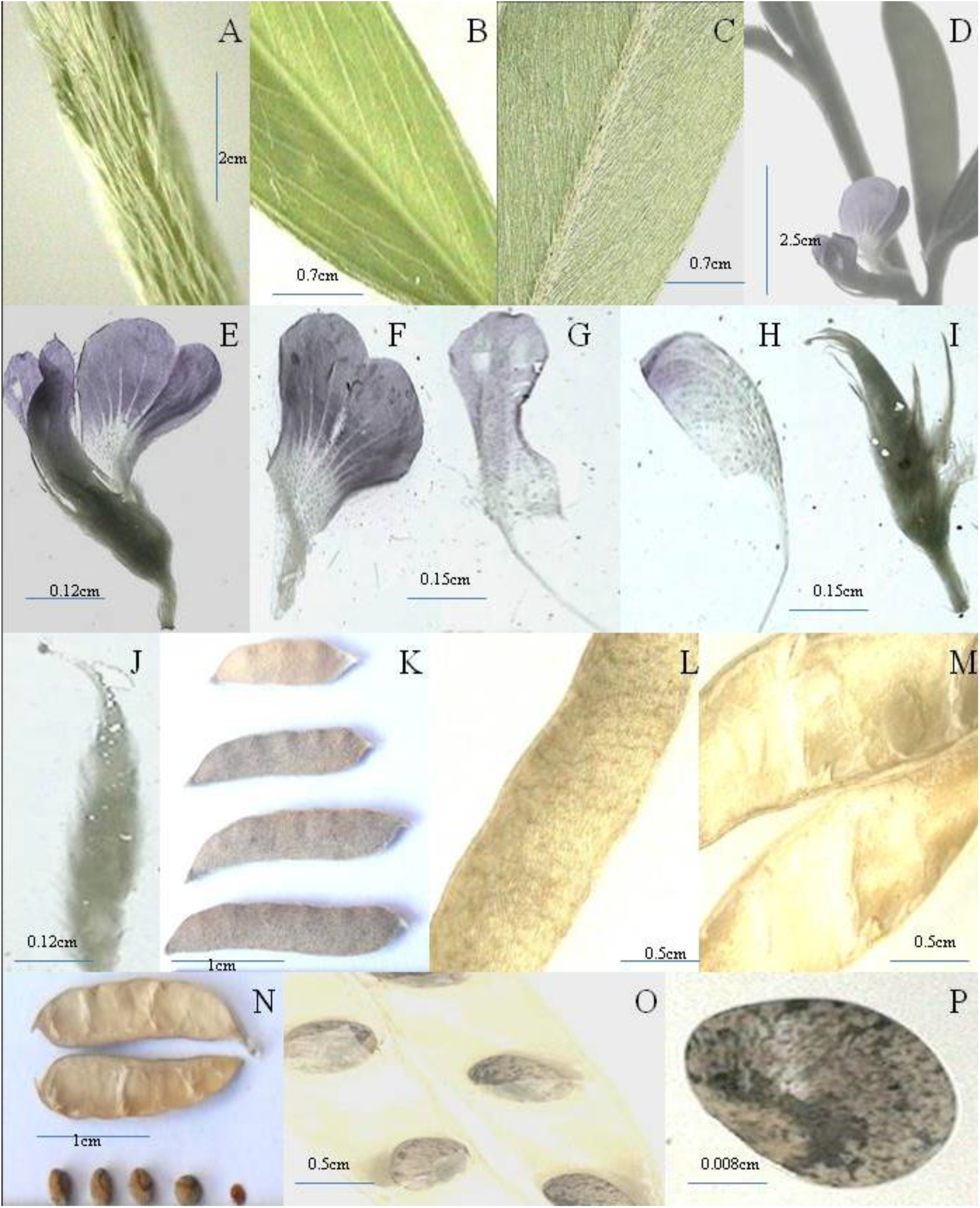
Detail Morphology Characters of *Tephrosia jamnagarensis*: A-stem, B- adaxial leaves portion, C- abaxial leaves portion D- inflorescence E- flower, F- standard, G- wing, H- keel, I- stamina sheath, J- Gynoceium, K- different size of pods, L- outer surface of pods, M- inner surface of pod N- pod and seed, O- seed arrangement in pod, P- seed.

### Plant morphology

***Tephrosia jamnagarensis*** syn. *Tephrosia axillaris* A.R.Sm. Proc. Natl. Inst. Sci. India, Pt. B, Biol. Sci. 24: 133. 1958. Vs ***Tephrosia collina* var *lanuginocarpa*** V.S. Sharma syn. *Tephrosia collina* V. S. Sharma. Journal of the Bombay Natural History Society 60: 758. 1963.

### Description

**Habit:** Annual herb with height of about 60-82 cm vs (Fig.4.I) vs Annual erect herb with height of about 60-100 cm (Fig.5.I). **Stem:** Simple or sparsely branched (Fig.4.G) covered with densely appressed silky hairs, erect or sub erect cylindrical vs Simple or quite above the base, rarely suffrutescent, terete and thinly pubescent towards base, angled or occasionally subterete, sometime villous and brown/pink striped above, at length sub glabrous (Fig.5.G). **Leaves:** Simple, 3.0-6.4 cm long, 6-9.5 mm board, stipules subulate, up to 3 mm long, uninerved, pubescent on margin; shortly petiolate up to 2-4 mm with silky appressed hair; lamina oblong –linear or elliptic linear, rounded to attenuate at the base, subacute-obtuse at apex, rarely mucronate, adaxially sub glabrous (Fig.1.B) and silky villous abaxially (Fig.1.C), lateral nerve 25-30, margin entire, reticulate venation with alternate phyllotaxy vs leaves compound imparipinnate (Fig.4.F); stipules up to 8mm long, subulate, 3 nerved persistent, leaf-rachis up to 16 cm long. abaxially furrowed, obscurely pulvinate at base; petiolate 1-2.5 cm long, tapering toward apex; leaflets 2-6.5 cm long, 0.5-1.7 cm broad, terminal are usually longer than laterals; 9-11 pairs, opposite or a few casually alternate, the last leaflets shift closure to main axis its distance from other leaflet is 1.3-2.17 cm while that from main axis is 0.2-1.5 cm, lamina of each leaflets is oblong – elliptic, 3-4.5 cm long, 6-9 mm board, rounded to attenuate at the base, obtuse or somewhat truncately emarginate at apex, rarely mucronate, sericeous abaxially and adaxially glabrous; margin entire, reticulate venation. **Inflorescence:** Axillary cyme (flower single or in pairs in rare case in group of three (Fig.1.D) vs 10-30 cm long, lax terminal racemes, much elongate, pedunculate, 5-9 flowered from usually above the middle or still higher up, 1-2 flowers at each node very rarely at the lowermost node subtended by a small leaf (bract); bracts are smaller than the pedicels, deciduous, similar to stipules (Fig.5.H). **Flower:** Mauve, 6.5mm long, shortly pedicile (2-3 mm), bracteate, ebracteolate, pentamerous, zygomorphic, hermaphrodite (Fig.1.E). vs flowers creamy white-sometime with light pink tinch, 1.5-1.7 cm long, bract filiform, bracteolate, shortly pedicile 1.5-2.5 mm densely argenteo-hirsue, zygomorphic, hermaphrodite, pentamerous (Fig.5.H). **Calyx:** 3.5 mm long, gamosepalous, valvate, campanulate with unequal lobes with silky appressed hairs, persistent in fruits (Fig.1.I). vs calyx 4-5 mm wide, 1.5-3 mm long, gamosepalous, campanulate with unequal lobes appressed with hairs, acuminate, persistent (Fig.2.B). **Corolla:** Polypetalous, vexillary aestivation with Standard, wing and keel. **Standard (vexllium)-** 4.5mm –broadly obovate to cordate, emarginate at the apex, punctate hairy toward outer side inner surface is glabrous, retuse at base (Fig.1.F); **Wing (alae)-** 5.5 mm – long, oblong, auriculate above the claw, punctate base, pubescent (Fig.1.G); **Keel (carina)**-3.4 mm long, auriculate, punctate, the 2 keel are joined atapical portion covering the staminal tube, mostly glabrous, and retuse at base (Fig.1.H); the claws of the wings and keel as long as their laminae vs polypetalous, Vexillary aestivation with Standard, wing and keel vs **Standard (vexillum)** 1.5 cm long, 1.2 cm broad, obovate, punctate, unguiculate, abaxially adpressed with silky brown hairs denser and longer in middle, margin ciliolate (Fig.2.C); **Wing (alae)** 1.4 cm long 4 mm wide, oblong, punctate, eared above claw (Fig.2.E); **Keel (carina)** −1.5 cm long, 5mm wide, punctate, glabrous, apex retuse with outer straight margin (Fig.2.F). **Androceium:** 4 mm long stamina sheath; anther 0.2 mm. long 10 in 9+1 dialdephous conditions, anthers all fertile, dithecous, introse (Fig.1.I). vs 1.2 cm long 3 mm broad staminal sheath; filament 2.5-3.5 mm long; anther 0.5mm long 0.2 mm wide 10 in 9+1 diadelphous, fertile, ditichous, introse (Fig.2.F). **Gynoceium:** 4 mm long, ovary subsessile, densely pubescent, monocarpellary, syncarpous, unilocular, superior with marginal placentation, style-short glabrous; stigma capitate (Fig.1.J) vs Gynocium 1.3 mm long, ovary sessile, densely pubescent, monocarpellary, unilocular, syncarpous, superior ovary with marginal placentation, style-2.5 mm long, glabrous; stigma minute, glabrous (Fig2.G). **Legume:** 2 X 0.5 cm long, densely and patently hairy with grayish tinge, oblique at both ends, apiculate, number of seed in each pod is 2-6 (Fig1.K.). vs legumes light brown, 6.5-8 cm long, 0.6-0.7 broad compressed, 8-13 seeded, straight and slightly falcate towards apex, sutures is conspicuously fringed with dull-brown, stiff and almost erect short hairs of uniform length, values dehiscence twisting completely by 3-4 turns (Fig.2.H). **Seed:** 1-3 mm long, 1.5-3 mm broad, roundish elliptical, muddish brown in color with black ornamentation near hilum on the seed coat. The hilum is circular and black in color at the level of seed coat. The seed coat is smooth and shiny in the texture. (Fig1.N) vs Blackish brown in color with black ornamentation encircling the hilum on the seed coat. The hilum is sub center positioned on the seed and black in color. The shape of the hilum was circular and in the slight depression on the seed coat. The texture of the seed coat is rough and non-shiny. The shape of the seed is oblong with 3mm-5mm in length to 2mm −3.5 mm in breadth (Fig.2.K).

**Fig.2.**
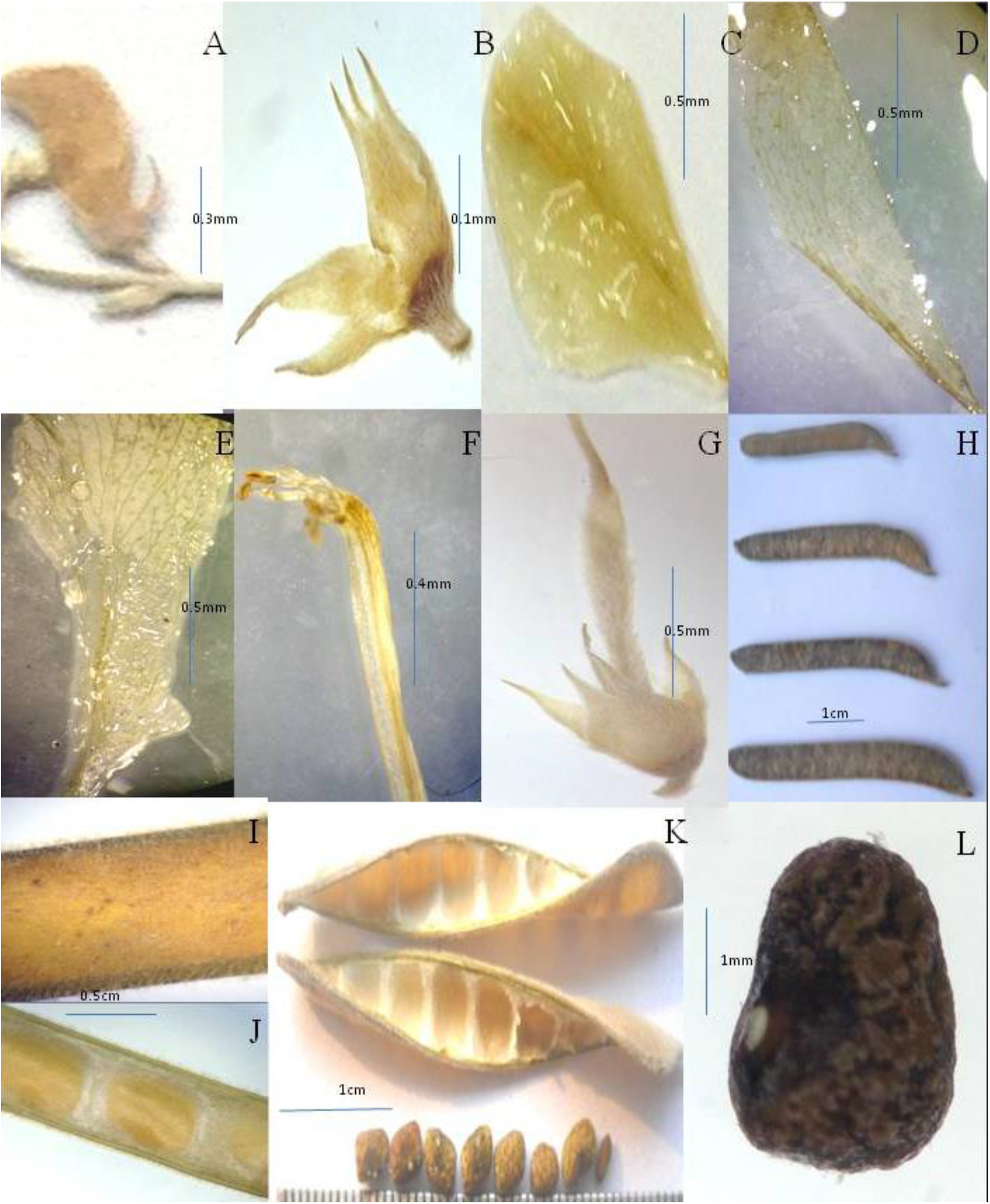
Detail Morphology Characters of *Tephrosia collina var lanuginocarpa*: A- flower, B- calyx, C- standard, D- flower, E- keel, F- staminal sheath, G- gynoceium, H- different size of pods, I- outer surface of pod, J- inner surface of pod, K- pod and seefs, L- seed

**Fig.3.**
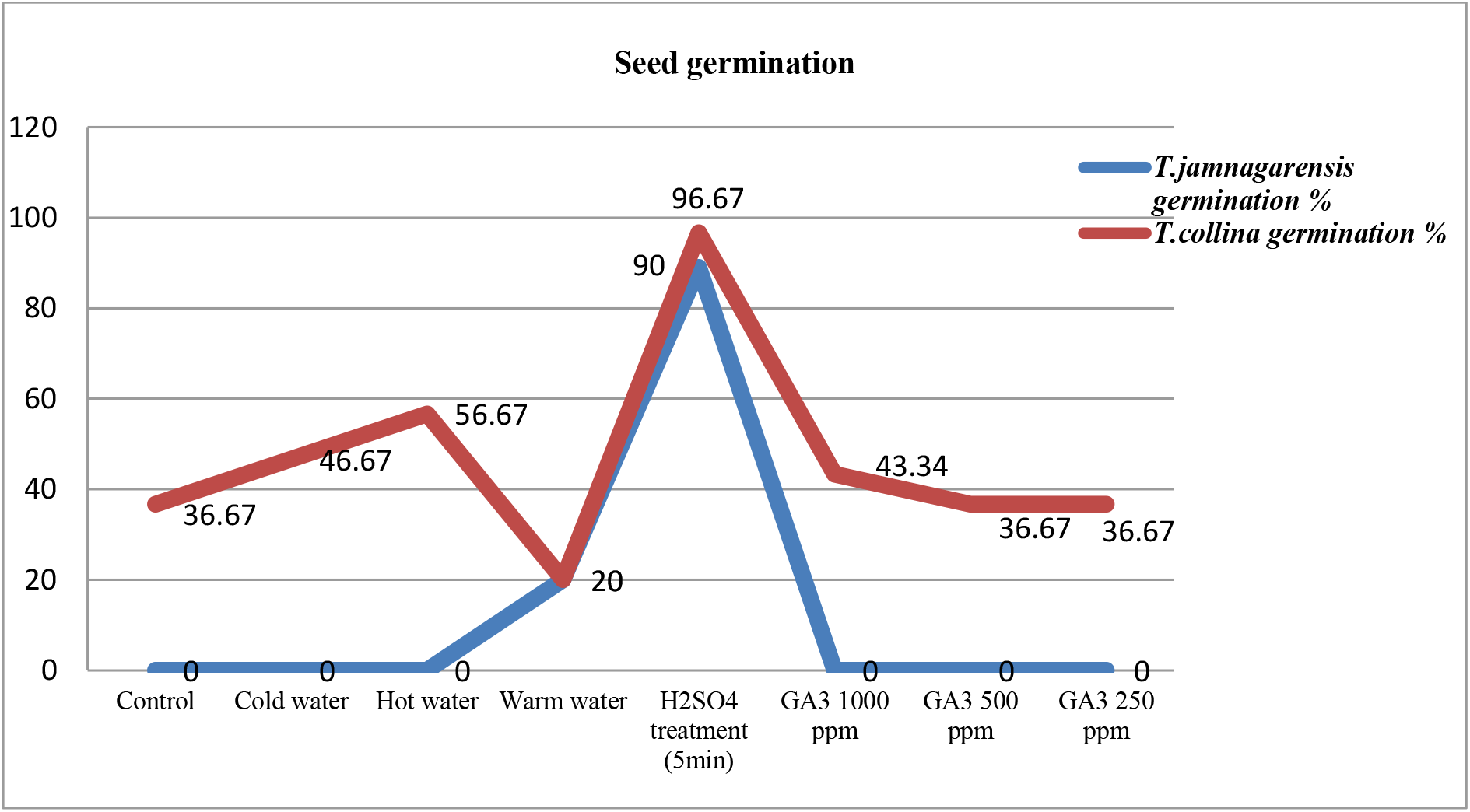
comparison between germination percentage and seed treatment of endemic species

**Fig 4.**
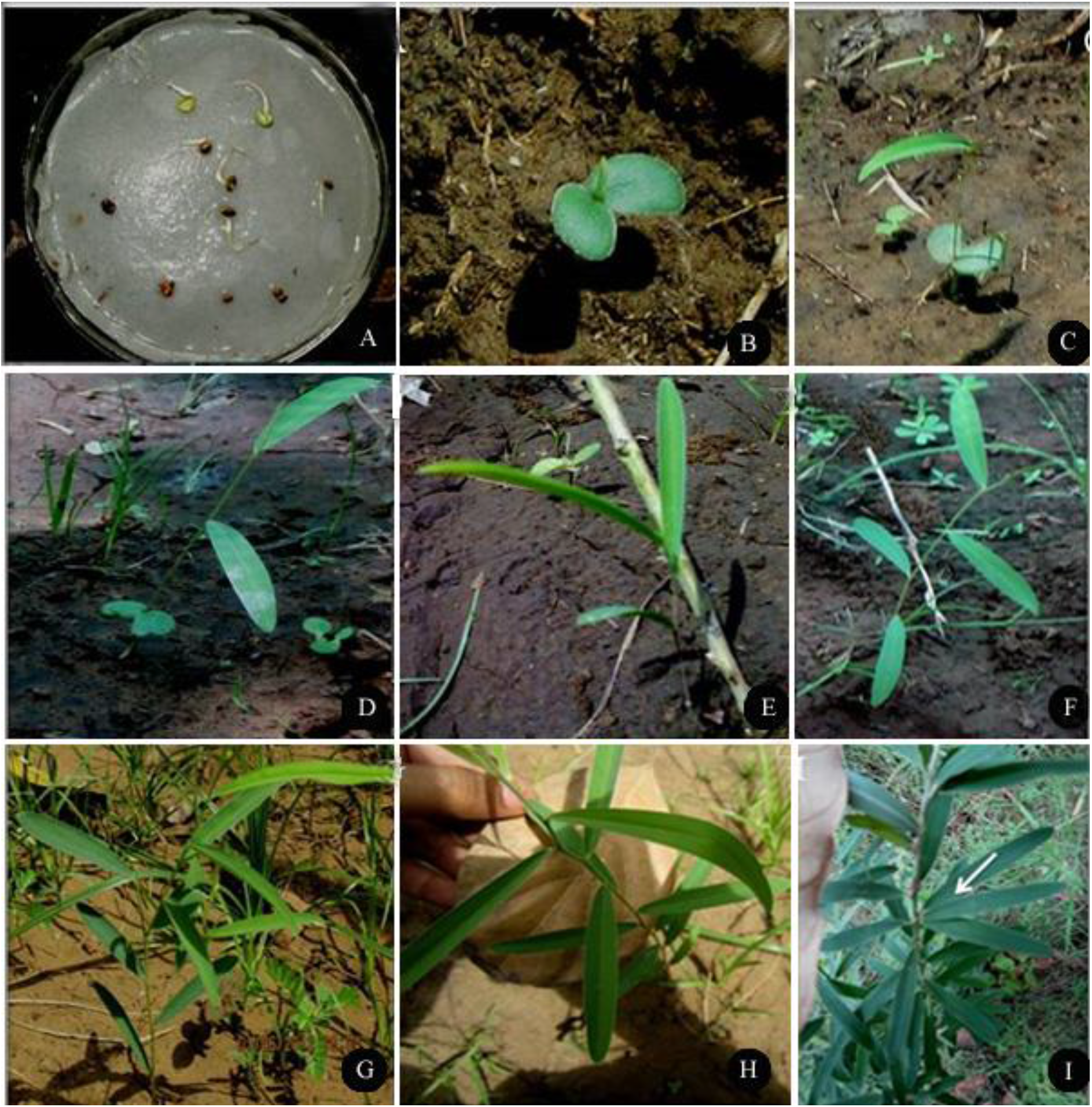
A. Germinating seed, B- fist stage leaf development, C- second stage leaf development, D- Third stage leaf development, E- fourth stage leaf development, F- sixth stage leaf development, G- branching, H- flower buds, I- Mature plant.

**Fig 5.**
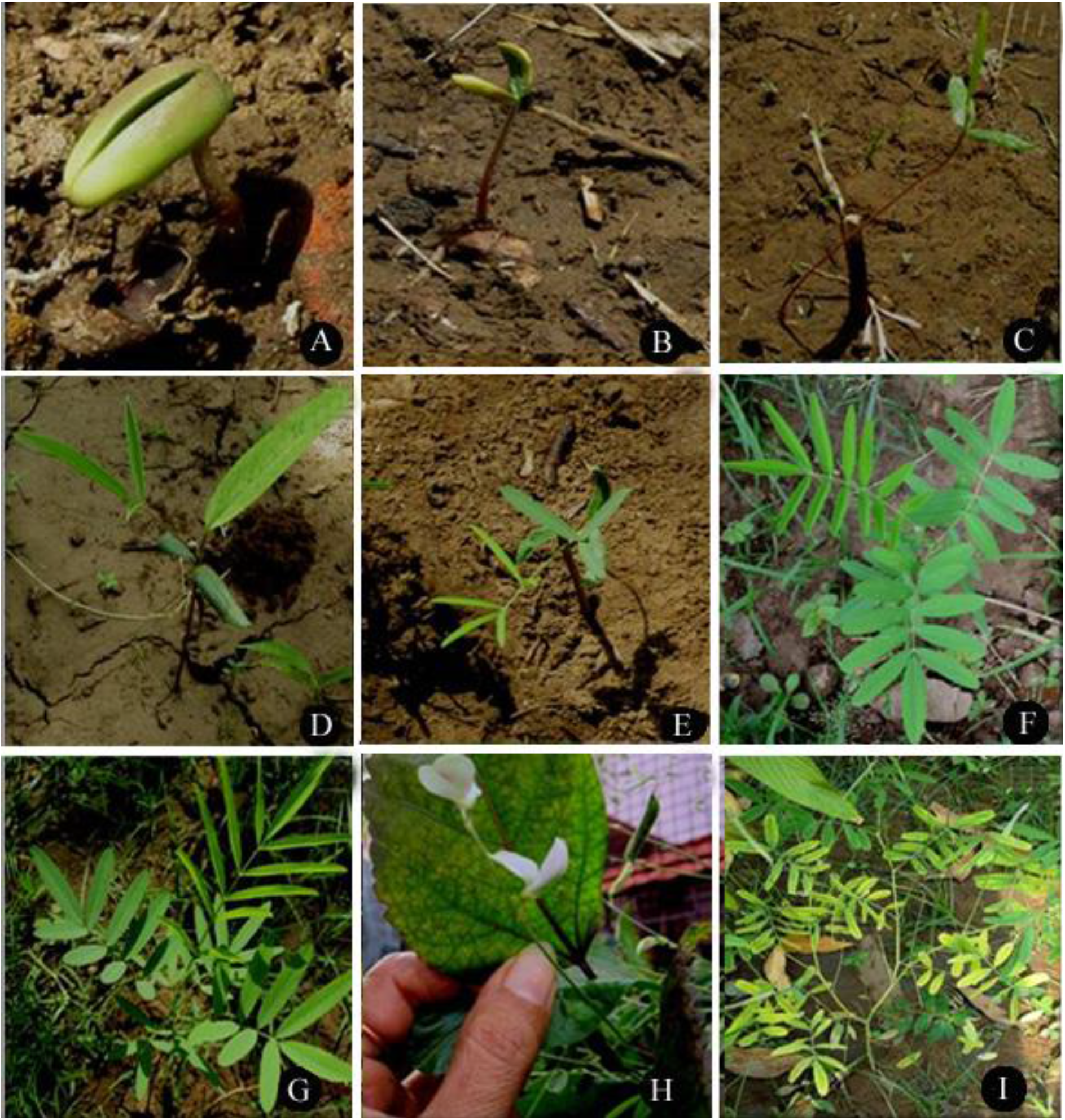
A- spouted seed, B- first stage leaf development, C- second stage leaf development, D- third stage leaf development, E- fourth stage leaf development, F- sixth stage leaf development, G-branching, H- flowering and fruiting stage, I- mature plant.

**Fig. 6:**
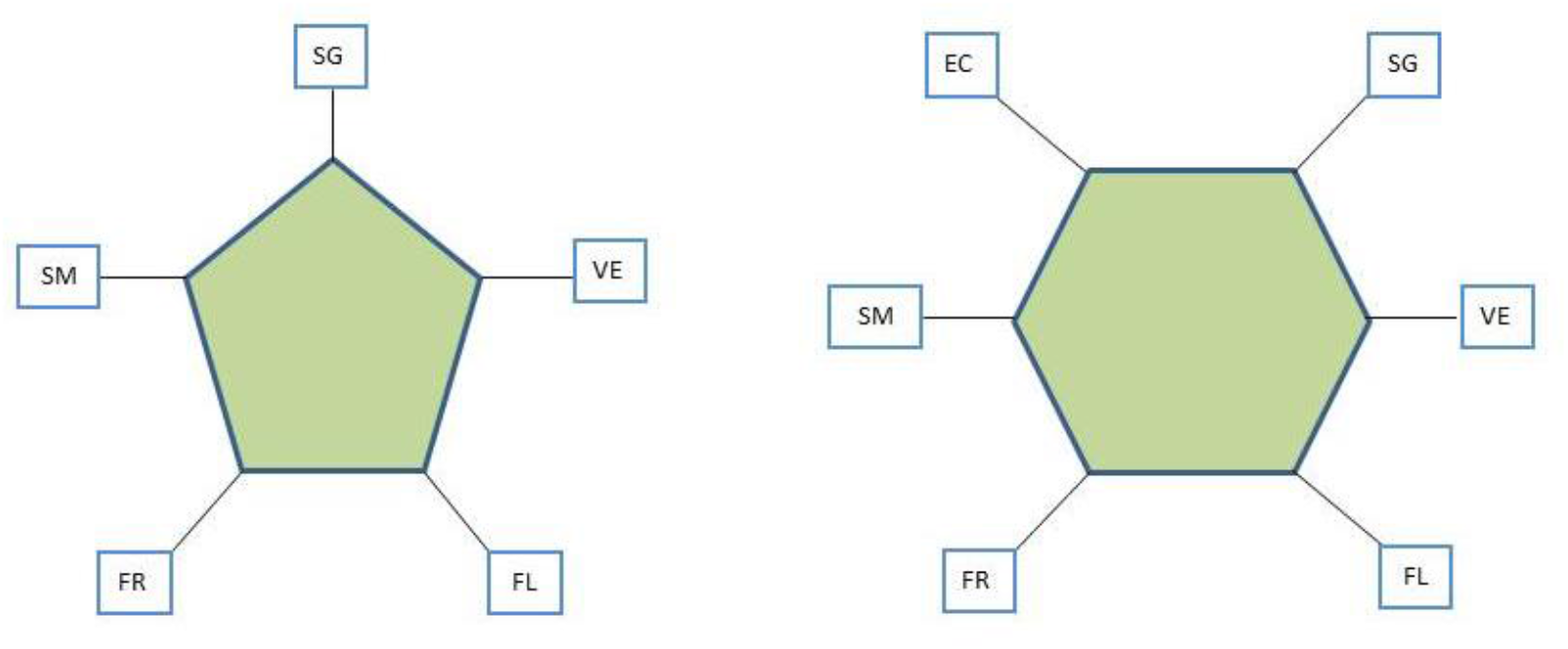
SG- Seed germination, VE- vegetative stage, FL- flowering period, FR- fruiting period, SM- seed maturation, EC- end of life cycle.

### Seed germination & phenology

Both these plant species were treated with various treatment methods of seed germination. Detailed experimental studies for seed germination in *T. jamnagarensis* and *T. collina var. lanuginocarpa* were summarizes in Table 2. The viability of *T. jamnagarensis* seeds is 12.5%, while that of the *T. collina var. lanuginocarpa* seed is 66.67%. It was observed that H2SO4 treatment for five minutes was the most effective treatment, showing 90% germination of seed, the other treatments such as cold water and different concentrations of GA3 (250, 500, 1000 ppm) tended to be moderately effective and Seed germination with warm water is the least effective.

**Table 2:** Seed germination in *T. jamnagarensis* and *T. collina* var *lanuginocarpa*

| Seed treatment of *T. jamnagarensis* |  |  |  |  |  |  | Seed treatment of *T. collina* |  |  |  |  |  |  |  |  |  |  |
| --- | --- | --- | --- | --- | --- | --- | --- | --- | --- | --- | --- | --- | --- | --- | --- | --- | --- |
| Sr. No. | Seed treatment | Day 1 | Day 2 | Day 3-15 | Total | Germination % | Day 1 | Day 2 | Day 3 | Day 4 | Day 5 | Day 6 | Day 7 | Day 8-14 | Day 15 | Total | Germination % |
| 1 | Control | - | - | - | - | 0 | 8 ± 1.53 | - | - | 1 ± 0.58 | - | 1 ± 0.58 | - | - | 1 ± 0.58 | 11 ± 2.3 | 40 |
| 2 | Cold water | - | - | - | - | 0 | 6 | 2 ± 1.15 | - | 2 ± 0.58 | - | 1 | 1 ± 0.58 | - | 2 ± 0.58 | 14 ± 1.53 | 50 |
| 3 | Hot water | - | - | - | - | 0 | 13 ± 2.08 | 2 ± 0.58 | - | 2 ± 0.58 | - | - | - | - | - | 17 ± 1.53 | 60 |
| 4 | Warm water | 1 | - | - | - | 0 | 3 | - | - | 2 ± 0.58 | - | 1 ± 0.58 | - | - | - | 6 ± 1 | 20 |
| 5 | H<sub>2</sub>SO<sub>4</sub> treatment (5min) | 30 | 28 ± 2.88 | - | - | 90% | 29 ± 0.58 | - | - | - | - | - | - | - | - | 29 ± 0.58 | 96.67 |
| 6 | GA<sub>3</sub> 1000 ppm | - | - | - | - | 0 | 8 ± 1.15 | 1 ± 0.58 | - | 3 ± 1 | - | 1 ± 0.58 | - | - | - | 13 ± 0.58 | 43 |
| 7 | GA<sub>3</sub> 500 ppm | - | - | - | - | 0 | 7 ± 1.15 | - | - | 1 ± 0.58 | - | 2 ± 0.58 | 1 ± 0.58 | - | 1 ± 0.58 | 11 ± 3.05 | 40 |
| 8 | GA<sub>3</sub> 250 ppm | - | - | - | - | 0 | 10 ± 2.3 | - | - | 1 ± 0.58 | - | - | - | - | - | 11 ± 2.08 | 40 |

**Table 3.**
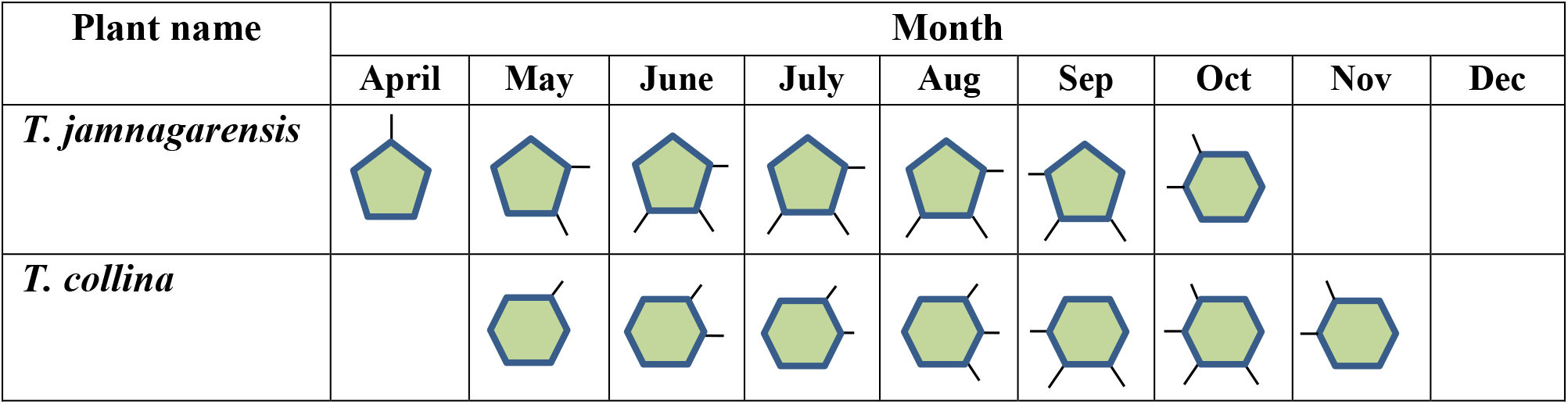
Comparative phenological characteristics of two endemic species.

The other species of the same genus *viz. T. purpurea*, which likewise germinates well when the seeds were treated with concentrated H_2_SO_4_ for five minutes Seeds of some species, including *T. bracteolata*, *T. candida*, *T. linearis*, and *T. vogelii*, can be effectively germinated by soaking in warm water for 45°C (Dharmalingam, Madhavrao, and Sundararaj 1973).

The seedlings of *T. jamnagarensis* were transplanted in the experimental site (Arboretum; The M. S. University of Vadodara, Gujarat.) in month of April. Vegetative growth continues until August. The flowering and fruiting of the plant begin in the month of May and continue until the end of July, plant complete its life cycle in the month of September.

Similarly, in *T. collina* seeds were transplanted to the same site in the month of May. Vegetative growth was observed as to be continues until August. Flowers began to bloom at the end of August. During September and October; the plant was at its peak in terms of flowering. Toward the end of September; fruit formation started, and it continues until the end of November and plant completes its annual life cycle, by this stage, plants have turned yellow and started dropping leaves

### Anatomy

#### Root Anatomy

Transverse view *T. jamnagarensis* vs *T. collina* var *lanuginocarpa* (fig.7); shows a circular thick outline with diameters ranging from 0.45-0.54 mm vs 1-10 mm (fig.7. A1, A2). The outermost layer in *T. jamnagarensis* consists of 7-10 layered cork cells with narrow and composed of 7-8 layers of parenchyma cells vs cork cells in *T. collina* var *lanuginocarpa* 0.19-0.36 mm composed of small squares cell which are 31-36 X 67-79 μm (fig.7 B1, C1). The ground tissues in *T. collina* var *lanuginocarpa* consist of four-five layer of parenchyma 81.3-114 X 53-59 μm (fig.7 B2, C2). The cortex of *T. jamnagarensis* 25-85 μm which is interrupted by phloem, and consists of prominent fiber patches 25-62.5 μm vs in *T. collina* var *lanuginocarpa* phloem 36-94 X 46-127 μm which is composed of sieve tube and phloem fibre. Xylem tissue in *T. jamnagarensis* contains rhomboidal calcium oxalate crystals which are composed of xylem fibers, xylem vessels and xylem parenchyma. Pith is absent (fig.7 D1, D2).vs secondary xylem occupied by secondary xylem which composed of xylem fibers, trachieds and xylem vessels which are 38-120 X 27-141μm size pith was not absurd.

**Fig. 7:**
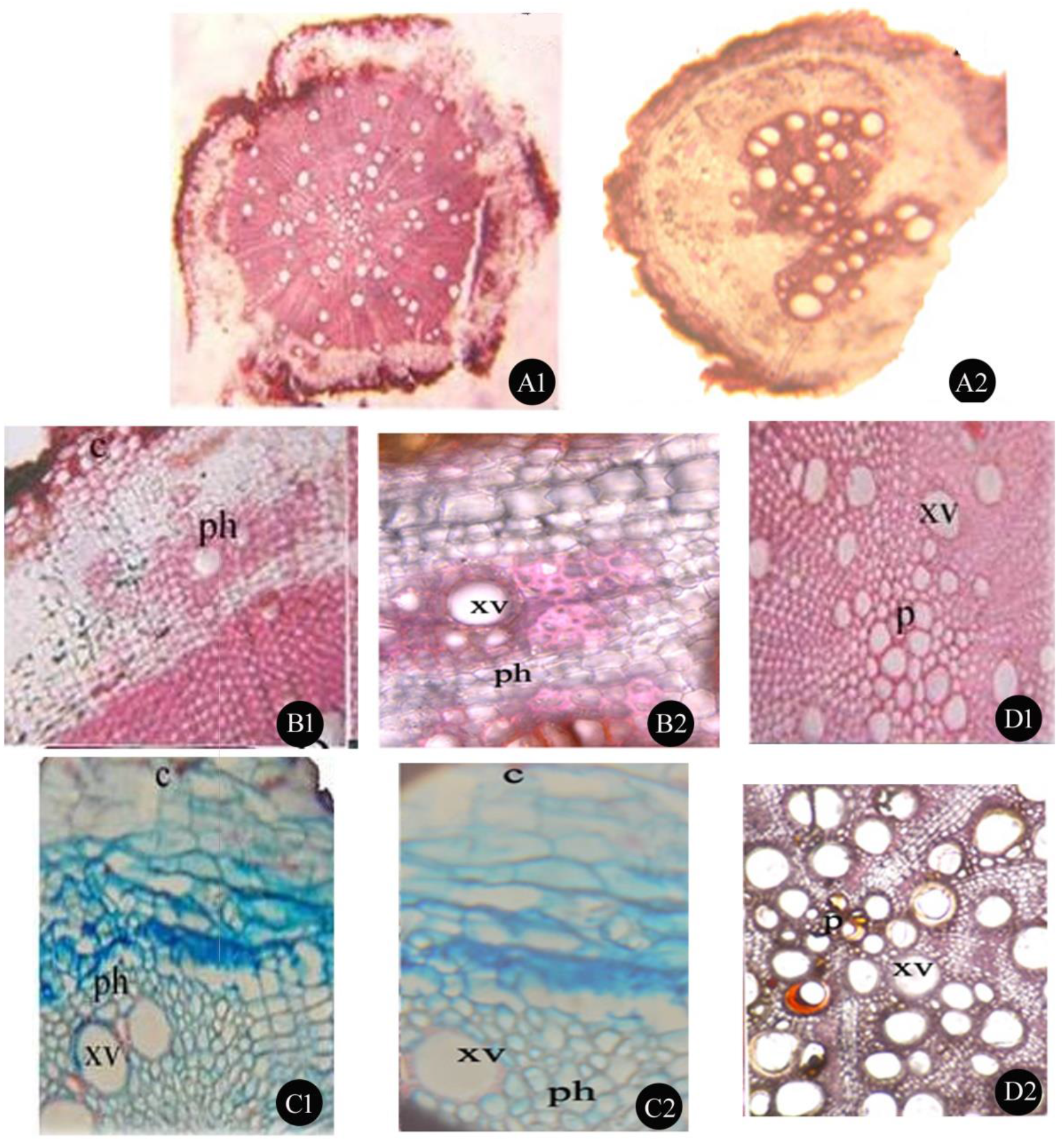
T.S. of Root: **A1:** T.S of root *T. jamnagarensis*, **A2**: T.S of root *T. collina* var *lanuginocarpa*, **B1, C1:** Cork region of *T. jamnagarensis* **B2, C2:** Cork region of *T. collina* var *lanuginocarpa*, **D1:** Pith region of *T. jamnagarensis*, **D2**: Pith region of *T. collina* var *lanuginocarpa*. **Legend**: Xv- xylem vessels, ph- phloem.

#### Stem anatomy

In fully grown mature stem of *T. jamnagarensis* have a circular outline with a diameter ranging from 0.81 to 1 mm when viewed transversely (Fig. 8 A1). Fig. 8 E shows barrel-shaped cells in *T. jamnagarensis* lined by a thin cuticle, and trichomes that are warty and unicellular 375 × 18.75 mm. The outer layer of *T. jamnagarensis* consists of two layers of collenchyma 27.5 mm thick followed by several layers of chlorenchyma 26.5 mm patches alternated with parenchyma patches 27.5 mm thick. Pericycle is composed of an interrupted ring of sclerenchyma patches containing calcium oxalate crystals. Endodermis barrel-shaped cells are found above the pericycle. Phloem is found in patches that are seprated by large medullar cells (Fig. 8 B1). Xylem is composed of angular vessels 23.8 mm, paedomorphic rays with calcium oxalates crystals and xylem fibers. Several patches of phloem are visible along the pith. The pith is large round and paranchymatous (Fig. 8 C1).

**Fig. 8:**
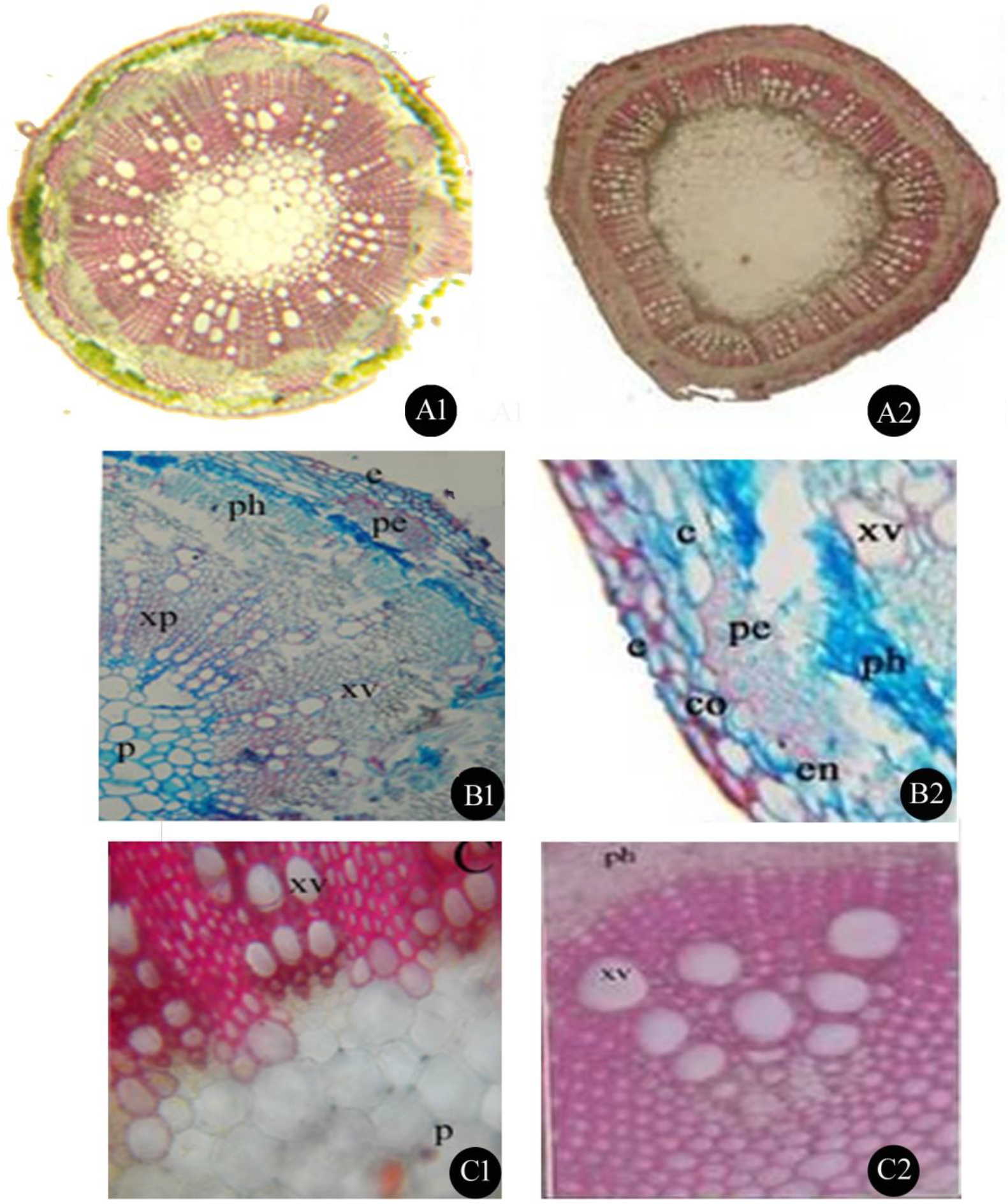
T.S. of Stem: **A1:** T.S of stem *T. jamnagarensis*, **A2**: T.S of stem *T. collina* var *lanuginocarpa*, **B1:** Enlarged view of stem *T. jamnagarensis* **B2:** Enlarged view of stem *T. collina* var *lanuginocarpa*, **C1:** Pith region of *T. jamnagarensis*, **C 2**: Pith region of *T. collina* var *lanuginocarpa*. **Legend**: e- epidermis, co- collenchyma, ch- chlorenchyma, en- endodermis, pe- pericycle, xv- xylem vessels, ph- phloem, p- pith, t- trichome

In transvers view of *T. collina* var *lanuginocarpa*, have an angular outline with a thickness of 1-10 mm (Fig. 8 A2). Epidermis composed of a single layer 15-29 X 12-60 mm with a lumen that is covered by thick cuticle. Hypodermis is composed of collenchyma 29-70 X 15-50 mm and chlorenchyma 117 X 80 mm patches. The endodermis is single layered, surrounded by sclerenchymatous pericycle, which are three-four layers deep. The pericycle is characterized by a presence of prime shape calcium oxalate crystals 28-49 mm. secondary phloem consist of three layers 21-44 X 23-45 mm encircling secondary xylem represented by sieve tube and phloem parenchyma (fig. 8 B2). Secondary xylem consists of medullary ray, xylem vessels 81.3 X 83.36 mm, and tracheid segment. Parenchymatous pith shows presence of starch grain, whereas the portion of the center is void (fig. 8 C2).

#### Leaf Anatomy

The transvers section of leaf in *T. jamnagarensis*; shows presence of upper epidermis, palisade, spongy and lower epidermis (Fig. 9-A). Both the epidermis composed of large barrel shaped cells covered by a thick cuticle and anisocytic stomata with unicellular trichomes 343.5 X 65μm (Fig. 9-B & C). Several large bundles of vascular tissues are seen along the midrib. The pericycle is found on either side of the large vascular bundles 28.8 μm and is composed of sclerenchyma cells. The pericycle contains rhomboidal calcium oxalate crystals. The vascular bundle consists of xylem and phloem. Mesophyll composed of palisade and closely packed spongy tissues.

**Fig. 9:**
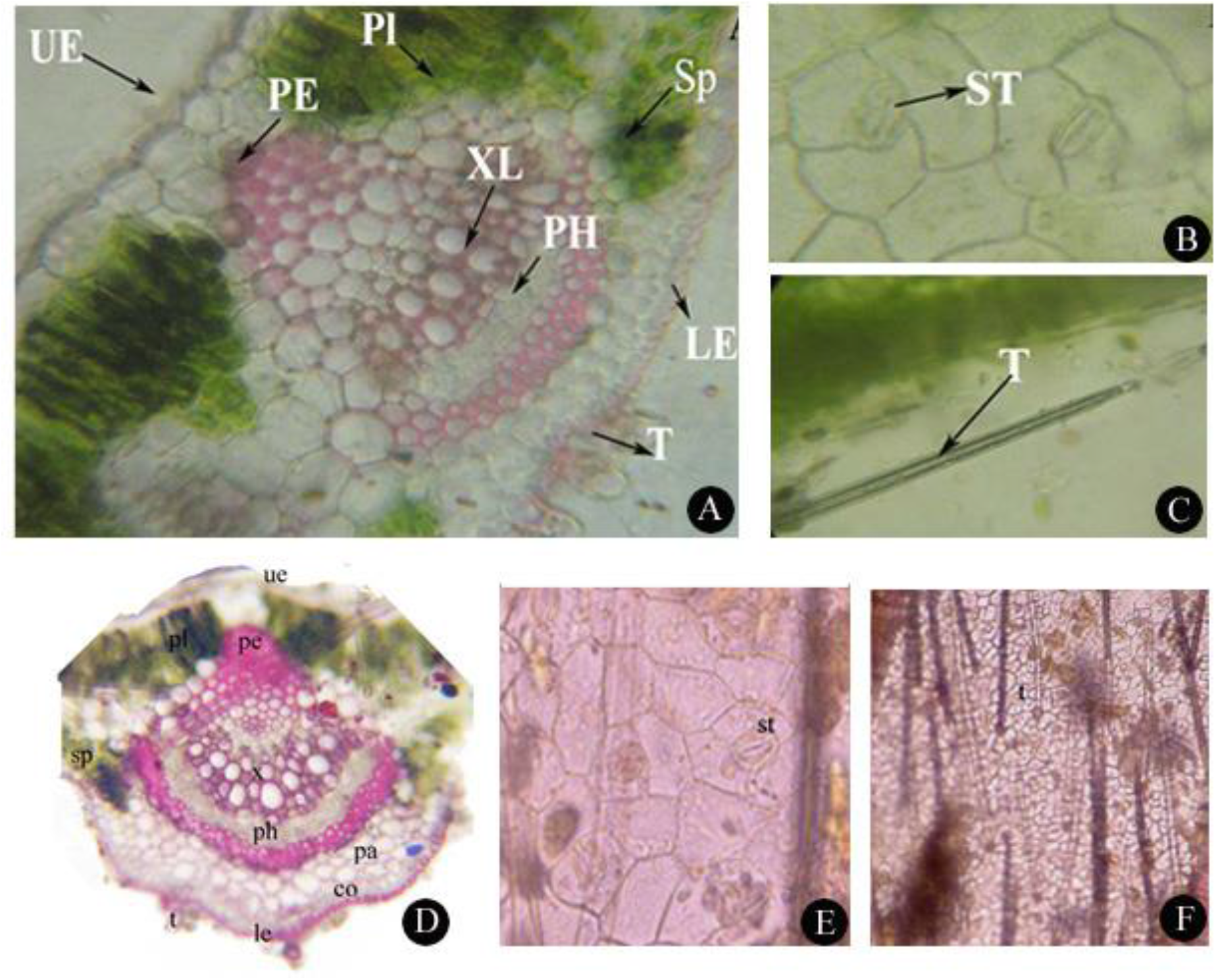
T.S. of Leaf: **A:** T.S of leaf *T. jamnagarensis*, **B**: anisocytic type of stomata *T. jamnagarensis*, **C:** Trichome of *T. jamnagarensis* **D:** T.S of leaf *T. collina* var *lanuginocarpa*, **E:** Stomata of *T. collina* var *lanuginocarpa*, **F**: Trichome of *T. collina* var *lanuginocarpa*. **Legend**: UE- upper epidermis, PL- pallisade, SP- spongy tissue, PE- pericycle, XL- xylem vessels, PH- phloem, P- pith, T- trichome, LE- lower epidermis, ST- stomata, T- trichome

In *T. collina* var *lanuginocarpa* the transvers section of leaves shows upper and lower epidermis 21-39X 18-35 mm attached to thick cuticle (13-28 mm) along with trichomes and stomata (Fig. 9-D). Anisocytic stomata are present along with unicellular trichomes 241-372 X 12-60μm. Palisade consist of elongated, linear rows of 2-3 cells, whereas, spongy tissues composed of three layers of loosely packed rounded cells. The midrib is composed of vascular bundles with sclerenchymatous pericycle 97-123μm. The xylem consists of 4-5 smaller vessels. The phloem is composed of few thin-walled cells.

The values of stomatal index, vein termination number, vein islet number, and palisade ratio are presented in Table no. 4.

**Table 4.** Micrometry studies of leaves in *T. jamnagarensisa* and *T. collina* var *lanuginocarpa*

| Micrometry studies of leaves |  |  |  |
| --- | --- | --- | --- |
| Sr. No. | Name | *T. jamnagarensis* | *T. collina* var *lanuginocarpa*, |
| 1 | Stomatal index | 18.4±3.13 | 14±0.02 |
| 2 | Palisade ratio | 4.4±0.5 | 3.57 ±0.83 |
| 3 | Vein islet no. | 3.6 | 2.95 |
| 4 | Vein termination no. | 1 or 2 | 1 or 2 |

#### A Phylogenetic Analysis of Sequenced Data

A comparative study on phylogenetic relationship was performed in *T. jamnagarensis, T. collina* var *lanuginocarpa* and with those of *Tephrosia* species which are naturally occurred in India. During the study a total of twelve sequence of rbcL gene and eleven sequence of matK gene were generated and submitted to the NCBI (National Center for Biotechnology Information). The accessions numbers of newly generated sequences are listed in table no. 5, along with sequence obtained from NCBI. The alignments of these sequences were carried out with the help of ClustalW software. The obtained alignment results were subsequently analyzed with RaxML software using 1000 bootstrap replicates and the GTR + I model. The differences at the species level are evident in fig no. (Figure number) 10 and 11. Wherein, different species of Indian *Tephrosia* were examined. Fig no. 10 demonstrates the phylogenetic relationship in contrast to rbcL gene, wherein *T. jamnagarensis* (AGV006) is separated with its sister clade *T. pumila* (AGV001) with 97 bootstrap supports, while *T. collina* var *lanuginocarpa* (AGV007) is separated from *T. purpurea* (AGV002) and *T. purpurea* subap. *apollinea* with 98 bootstrap support. Similarly, *T. candida* (AGV010), *T. uniflora* (AGV004) and *T. vogelii* (AGV011) are genetically close but separated with the sister clade and bootstrap support of 96, 96 and 99 respectively. Furthermore, *T. villosa* (AGV005) and *T. pentaphylla* (AGV012) are close to each other and were separated with bootstrap support of 95, while both species have a close sister clade with *T. calophylla* (AGV009) and bootstrap support was 98 respectively. *T. strigosa* (AGV003) is having clear separation with the other species in support of 100 bootstraps value. Figure no. 11 demonstrates the phylogenetic graph which was analyzed by matK gene. All the species in figure no. 11 are separated in similar position when compared with the rbcL gene with only difference of bootstrap support values. *T. jamnagarensis* (AGV006) and *T. pumila* (AGV001) were having bootstrap difference of 90. While *T. collina* var *lanuginocarpa* (AGV007), *T. purpurea* (AGV002) and *T. purpurea* subsp. *apollinea* were having 99 bootstrap differences. Similarly, *T. candida* (AGV010), *T. uniflora* (AGV004) and *T. vogelii* (AGV011) were having 94, 78 and 92 bootstrap support. In *T. villosa* (AGV005) and *T. pentaphylla* (AGV012) having 95, while both species have a sister clade with *T. calophylla* (AGV009) with bootstrap difference of 90 respectively. In *T. strigosa* (AGV003) it shows similar clade separation when compared with rbcL gene with bootstrap support of 100.

**Table.5.**
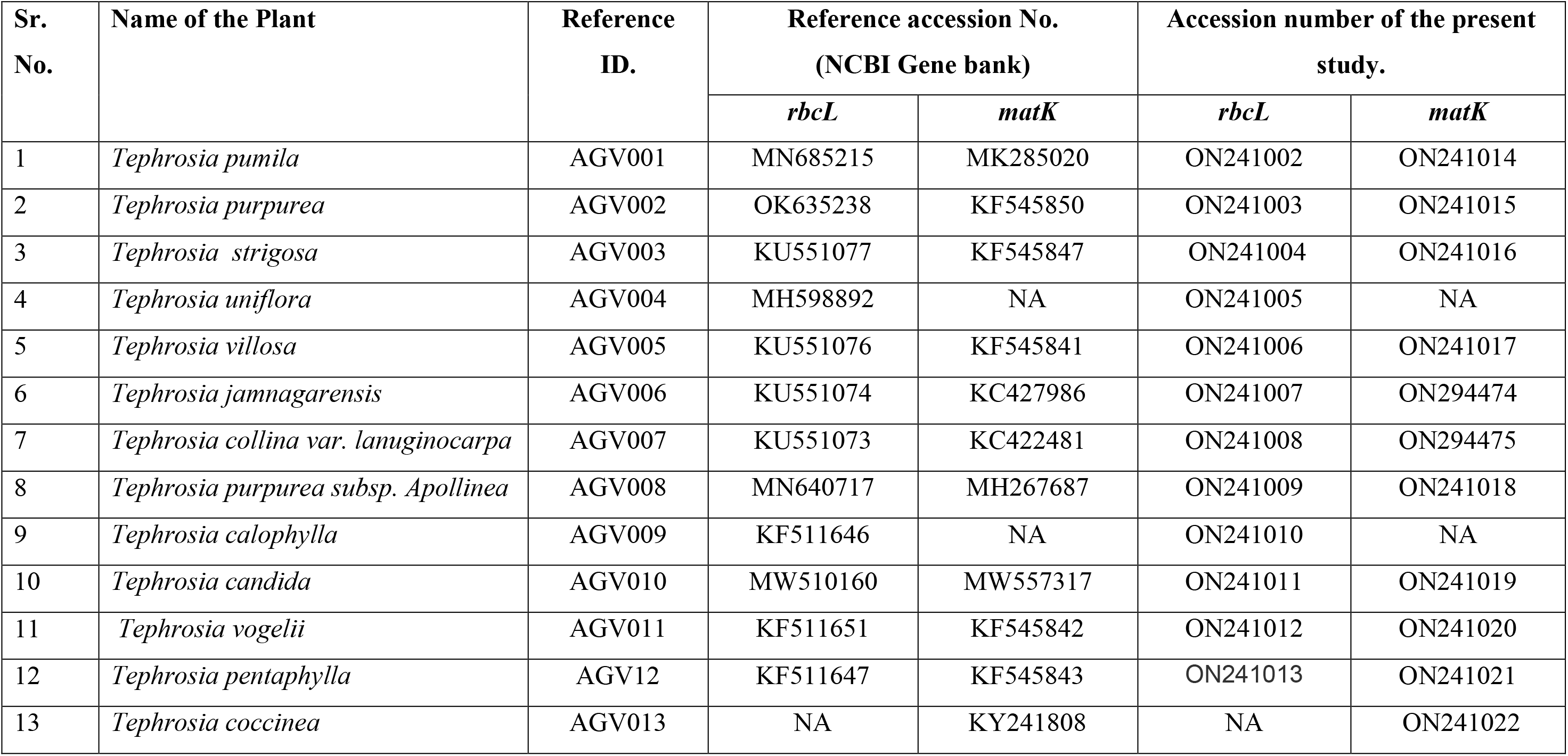
Accession no of reference sequence *vs* newly generated sequence.

| Sr. No. | Name of the Plant | Reference ID. | Reference accession No.<br>(NCBI Gene bank) |  | Accession number of the present study. |  |
| --- | --- | --- | --- | --- | --- | --- |
|  |  |  | *rbcL* | *matK* | *rbcL* | *matK* |
| 1 | *Tephrosia pumila* | AGV001 | MN685215 | MK285020 | ON241002 | ON241014 |
| 2 | *Tephrosia purpurea* | AGV002 | OK635238 | KF545850 | ON241003 | ON241015 |
| 3 | *Tephrosia strigosa* | AGV003 | KU551077 | KF545847 | ON241004 | ON241016 |
| 4 | *Tephrosia uniflora* | AGV004 | MH598892 | NA | ON241005 | NA |
| 5 | *Tephrosia villosa* | AGV005 | KU551076 | KF545841 | ON241006 | ON241017 |
| 6 | *Tephrosia jamnagarensis* | AGV006 | KU551074 | KC427986 | ON241007 | ON294474 |
| 7 | *Tephrosia collina* var. *lanuginocarpa* | AGV007 | KU551073 | KC422481 | ON241008 | ON294475 |
| 8 | *Tephrosia purpurea* subsp. *Apollinea* | AGV008 | MN640717 | MH267687 | ON241009 | ON241018 |
| 9 | *Tephrosia calophylla* | AGV009 | KF511646 | NA | ON241010 | NA |
| 10 | *Tephrosia candida* | AGV010 | MW510160 | MW557317 | ON241011 | ON241019 |
| 11 | *Tephrosia vogelii* | AGV011 | KF511651 | KF545842 | ON241012 | ON241020 |
| 12 | *Tephrosia pentaphylla* | AGV12 | KF511647 | KF545843 | ON241013 | ON241021 |
| 13 | *Tephrosia coccinea* | AGV013 | NA | KY241808 | NA | ON241022 |

**Fig 10.**
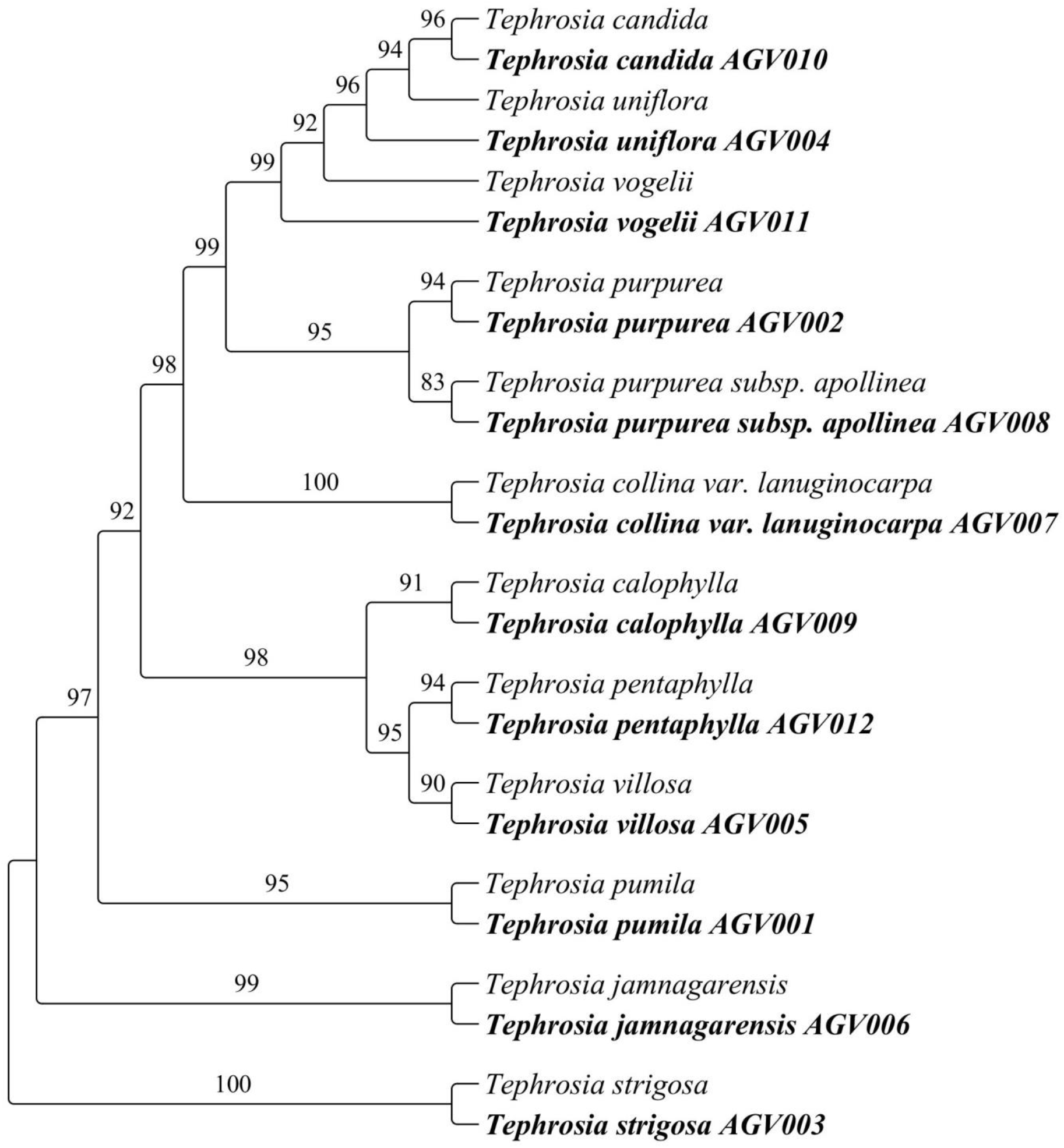
Phylogenetic graph of *T. jamnagarensis* & *T. collina*. using rbcL gene.

**Fig 11.**
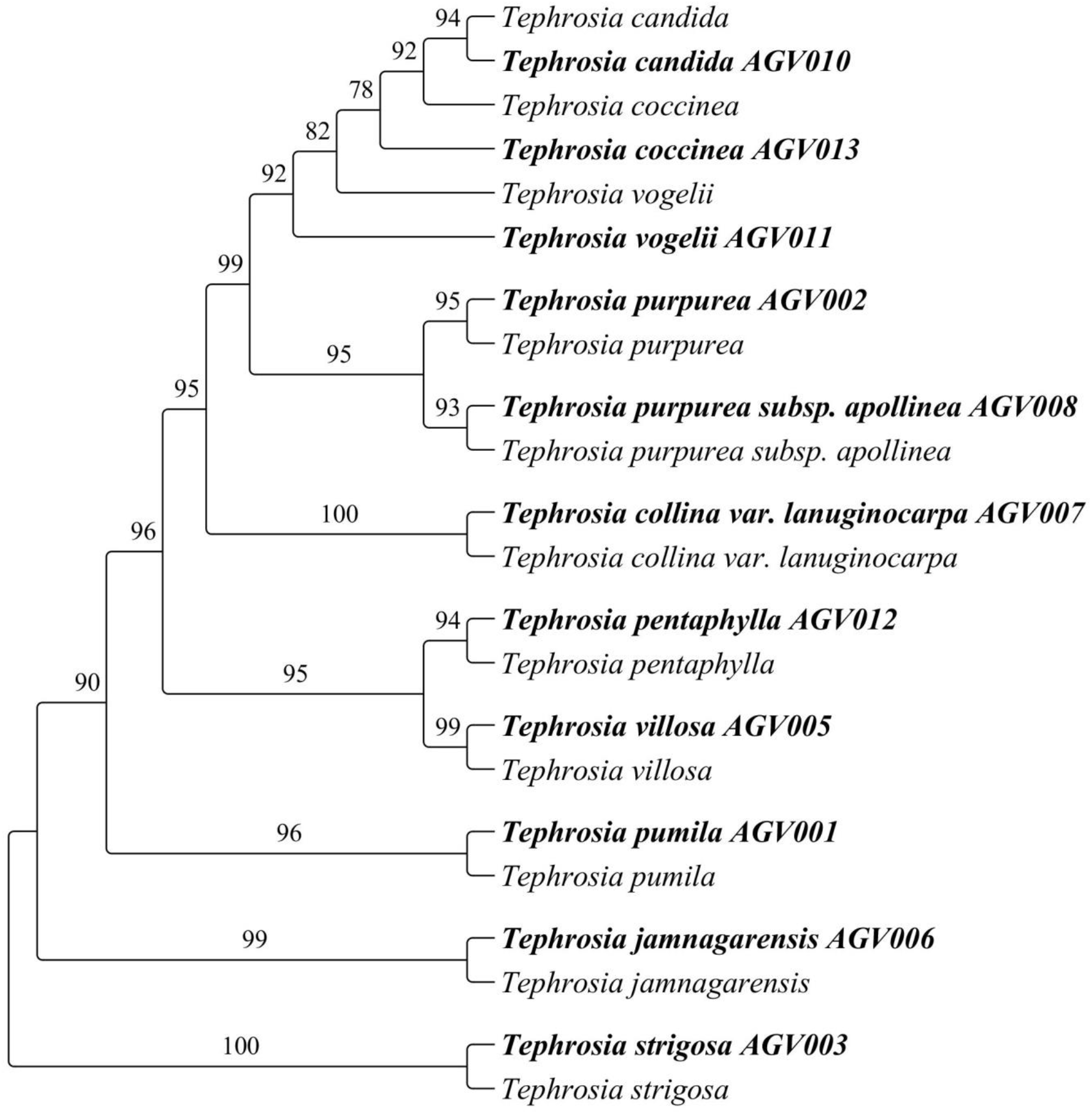
Phylogenetic graph of *T. jamnagarensis* & *T. collina*. using matK gene.

## Discussion

For the precise confirmation of any plant it requires morphology, anatomy and phylogeny investigation which plays an essential role to verifying the plant’s identity. These features help to verify taxa at a cryptic level which can be crucial for species-level identification. The genus is widespread throughout globe with highest number of species found in Africa-Madagascar (c.170 species), Australia (c. 90 species) and central and tropical North America (c. 45 species), Central and North America (c. 45 species) (G. Lewis et al. 2005) A total of 29 species and two subspecies one variety represented by South Asia (Kumar and Sane 2003) *T. jamnagarensis* was identified and reported first by Santapau (Santapau n.d.) from Gujarat. After to 3 decades Nagar et al. (Nagar, Sata, and Pathak 2003) reported as to be rediscovery from the type location; wherein detailed description of plant viz. morphology, habit, habitat, distribution and photography was provided. In 2016, *T. jamnagarensis* was assessed and listed under endangered category with B1 ab (i,ii,iv) + 2ab (I,ii,iv) for the IUCN red list data base (LIST IUCN 2017).

## Conclusion

Despite all the research to date, there has not been any study on the anatomical features, genetic discrimination or seed germination was reported so far. Therefore the aim of the study was to put together, for the first time ever, a complete comparison between *T. jamnagarensis* and *T. collina* var *lanuginocarpa* in terms of all essential features. This may prove critical for identification and conservation of both the species. Thus results of this study may be useful for exploratory research in the future.

## Acknowledgment

We are highly thankful to Head department of Botany, The M.S.University of Baroda, Gujarat for providing research lab facilities. Authors would like to extend their thanks to Mr. Ravi S. Patel, Ms. Amisha Patel and Nimisha Patel for necessary assistance. Authors also thanks to Shri. Govind Vanzara, Mr. Pradeep Vanzara, Mr. Amit Shrivastav, Mr. Kapil Yadav for their support and encouragement.

## References

Berlyn, Graeme P. 1976. Botanical Microtechnique and Cytochemistry.

Bole J.M., P V and Pathak. 1988. “Flora of Saurashtra (Pt. II & III).” Flora of India, ser. 2. Botanical Survey of India, Culcutta.

Chen, Yinning et al. 2014. “Natural Products from the Genus Tephrosia.” Molecules 19(2): 1432–58.

Cook, T. “The Flora of the Presidency of Bombay,.” Botanical survey of india, Hawrah 1: 611.

Dagne, Ermias, Abiy Yenesew, and Peter G Waterman. 1989. “Flavonoids and Isoflavonoids from Tephrosia Fulvinervis and Tephrosia Pentaphylla.” Phytochemistry 28(11): 3207–10.

Dharmalingam, C, S Madhavrao, and D Sundararaj. 1973. “Pregermination Treatment of Testing Seeds (Tephrosiapurpures Pers.) to Improve Germination.” Seeds research 1: 58–62.

Hall, Tom. 1999. “BioEdit: A User-Friendly Biological Sequence Alignment Editor and Analysis Program for Windows 95/98/NT.” In Nucleic Acids Symp. Ser.,, 95–98.

Kaplan, Donald R. 2001. “The Science of Plant Morphology: Definition, History, and Role in Modern Biology.” American Journal of Botany 88(10): 1711–41.

Karthikeyan Sanjappa, M. & Moorthy, S., S. 2009. “Flowering Plants of India-Dicotyledon.” Botanical survey of india, Kolkata.

Kumar, Sudershan, and P V Sane. 2003. Legumes of South Asia. Royal Botanic Gardens.

Lakshmi, P et al. 2008. “Genetic Relationship among Tephrosia Species as Revealed by RAPD Analysis.” Asian Journal of Biological Science 1(1): 1–10.

Lanfear, Robert, Brett Calcott, Simon Y W Ho, and Stephane Guindon. 2012. “PartitionFinder: Combined Selection of Partitioning Schemes and Substitution Models for Phylogenetic Analyses.” Molecular biology and evolution 29(6): 1695–1701.

Lewis, G, B Schrire, B Mackinder, and M Lock. 2005. “Legumes of the World, Royal Botanic Gardens, Kew.” Edinb. J. Bot 62: 195–96.

Lewis, Gwilym Peter, Brian Schrire, Barbara Mackinder, and Mike Lock. 2005. Legumes of the World. Royal Botanic Gardens Kew.

LIST, IUCN R E D. 2017. “Tephrosia Jamnagarensis. The IUCN Red List of Threatened Species 2017.” The IUCN Red List of Threatened Species 2017–3. https://www.iucnredlist.org/species/96238744/96239894.

Nagar, P S, S J Sata, and S J Pathak. 2003. “REDISCOVERY OF TEPHROSIA JAMNAGARENSIS (FABACEAE), AN ENDANGERED AND NARROW ENDEMIC PLANT SPECIES OF SAURASHTRA, GUJARAT, INDIA.” SIDA, Contributions to Botany: 1701–5.

Phillips, B G. 1986. “Webster Third New International of the English Language.” MerrianWebster Incorporation U. S.: 1, 44.

Quattrocchi, Umberto. 2017. CRC World Dictionary of Plant Names: Common Names, Scientific Names, Eponyms, Synonyms, and Etymology. Routledge.

Samuel, Vimal John, Agasa Ramu Mahesh, and Vedigounder Murugan. 2019. “Phytochemical and Pharmacological Aspects of Tephrosia Genus: A Brief Review.” J. Appl. Pharm. Sci 9: 117–25.

Sanjappa, M. 1992. Legumes of India. Bishen Singh Mahendra Pal Singh.

Santapau, H. “1958. Addition and Corrections to the Indo-Nepalese Flora.” Proc. Natl. Sei. Inst. India B 24.

Shah, Gopalkrishna Laljibhai. 1978. “Flora of Gujarat State.”

Silvestro, Daniele, and Ingo Michalak. 2012. “RaxmlGUI: A Graphical Front-End for RAxML.” Organisms Diversity & Evolution 12(4): 335–37.

Stuessy, Tod F, Daniel J Crawford, and Clodomiro Marticorena. 1990. “Patterns of Phylogeny in the Endemic Vascular Flora of the Juan Fernandez Islands, Chile.” Systematic Botany: 338–46.

Tamura, Koichiro et al. 2013. “MEGA6: Molecular Evolutionary Genetics Analysis Version 6.0.” Molecular biology and evolution 30(12): 2725–29.

Thaker, J I. 1910. “Vanasptai Sastra-Barda Dungar Ni Jadibuti Tani Pariksha Upyog. (Botany - A Complete and Comprehensive Account of the Flora of Barda Mountain (Kathiawad).” Gujarati Printing Press, Bombay: 717.

Thompson, Julie D, Toby J Gibson, and Des G Higgins. 2003. “Multiple Sequence Alignment Using ClustalW and ClustalX.”Current protocols in bioinformatics (1): 2.3. 1–2.3. 22.

Wood, Carroll E. 1949. “The American Barbistyled Species of Tephrosia (Leguminosae).” Contributions from the Gray Herbarium of Harvard University (170): 193–iv.

Yu, Jing, Jian-Hua Xue, and Shi-Liang Zhou. 2011. “New Universal MatK Primers for DNA Barcoding Angiosperms.”Journal of Systematics and Evolution 49(3): 176–81.

